# Protein Interaction Network-based Deep Learning Framework for Identifying Disease-Associated Human Proteins

**DOI:** 10.1101/2021.06.03.446973

**Authors:** Barnali Das, Pralay Mitra

**Affiliations:** Department of Computer Science and Engineering, Indian Institute of Technology Kharagpur, India - 721302

**Keywords:** Deep Learning-based Classification, Graph Convolutional Networks, Disease-associated Proteins, Topological features of Protein Locality Graph, Enrichment analysis

## Abstract

Infectious diseases in humans appear to be one of the most primary public health issues. Identification of novel disease-associated proteins will furnish an efficient recognition of the novel therapeutic targets. Here, we develop a Graph Convolutional Network (GCN)-based model called PINDeL to identify the disease-associated host proteins by integrating the human Protein Locality Graph and its corresponding topological features. Because of the amalgamation of GCN with the protein interaction network, PINDeL achieves the highest accuracy of 83.45% while AUROC and AUPRC values are 0.90 and 0.88, respectively. With high accuracy, recall, F1-score, specificity, AUROC, and AUPRC, PINDeL outperforms other existing machine-learning and deep-learning techniques for disease gene/protein identification in humans. Application of PINDeL on an independent dataset of 24320 proteins, which are not used for training, validation, or testing purposes, predicts 6448 new disease-protein associations of which we verify 3196 disease-proteins through experimental evidence like disease ontology, Gene Ontology, and KEGG pathway enrichment analyses. Our investigation informs that experimentally-verified 748 proteins are indeed responsible for pathogen-host protein interactions of which 22 disease-proteins share their association with multiple diseases such as cancer, aging, chem-dependency, pharmacogenomics, normal variation, infection, and immune-related diseases. This unique Graph Convolution Network-based prediction model is of utmost use in large-scale disease-protein association prediction and hence, will provide crucial insights on disease pathogenesis and will further aid in developing novel therapeutics.

## 1 Introduction

Human beings are susceptible to diseases caused by a variety of reasons including environmental factors, microorganisms (viruses and bacteria), and mostly, genetics. Since majority of the diseases in humans are caused by the genes, analyzing the genetic characteristics is a vital requirement for effective treatment of the diseases. Hence, deciphering the association between genetic diseases and their causal genes and proteins is a significant problem concerning human health. But this task is highly challenging due to the limited knowledge of diseases, genes, proteins, and their associations. Due to the pivotal roles played by the proteins in the cell, protein-protein interactions (PPIs) maneuver the biological processes responsible for the healthy and diseased states in organisms (Gonzalez and Kann, 2012). Research conducted in several studies derive a set of important observations highlighting the importance of Protein-Protein Interaction Networks (PPIN) in identifying unknown hereditary disease-associated proteins. It has been mentioned that disease pathways are already encoded in the human PPIN (Goh et al., 2007) and there is a tendency that the proteins involved in the same disease to share interactions in the PIN. Also, the number of interactions between the proteins associated with the same disease is 10 times higher than random expectation (Goh et al., 2007; Gandhi et al., 2006).

PPINs are capable of examining several biological processes in a systematic manner and numerous existing studies have already utilized and exploited the PINs for predicting potential disease-related proteins or drug targets. Examinations have been conducted on a variety of sequence features and functional similarities between known human hereditary disease genes and those not known to be involved with any disease. Based on the identified common patterns detected by these examinations, efficient classifiers have been designed by many studies. A good number of the existing research works apply machine learning-based techniques for determining the disease-associated proteins from the PIN of human. Protein sequence and protein interaction network features have been computed and integrated together into Support Vector Machine (SVM), Naive Bayes (NB), Random Forest (RF), and Deep Neural Network (DNN) models (Barman et al., 2019). Although the detailed architecture of the DNN is not available. Multiple combinations of these sequence and network features perform differently and different tests have been carried out to determine the combination delivering the best performance (Barman et al., 2019). Disease symptoms and the proteins’ primary sequence features characterize the diseases and the proteins, respectively (Chen et al., 2020). Again, the disease-gene associations are represented by grayscale images and a Convolutional Neural Network (CNN) model is trained utilizing these images which makes the method dependable on the image sizes and demands computational time accordingly (Chen et al., 2020). Masood et al. utilize the Dijkstra’s algorithm for computing the shortest paths between two known disease genes in the PIN in identifying the candidate disease genes (Masood et al., 2018). All the genes that are part of the all possible shortest paths between every pair of known disease genes are the potential disease-associated genes. An increase in the availability of information on the known disease-genes, increases the number of shortest paths to be calculated. Further, topological features of the human PIN have been exploited for training a KNN (Xu and Li, 2006) and a RF classifier (Yu et al., 2020).

Existing works for predicting the novel disease-associated proteins or genes require significant amount of data including human PIN, protein sequence features, PIN features, disease symptoms, and shortest path information to train their models (Barman et al., 2019; Chen et al., 2020). Mainly, topological analyses are performed on the PIN and machine learning-based techniques and image-based CNNs are exploited for this purpose. Here, we propose a novel Graph Convolutional Network (GCN)-based deep learning framework viz., *P*rotein *I*nteraction *N*etwork-based *De*ep *L*earning (PINDeL), through a systematic embedding of the Protein Locality Graph (PLG) and its topological features. PINDeL considers the protein spatial locality information in the form of protein locality graph of Homo sapiens (human) as constructed by the network-based zoning methodology (Das et al., 2019), alongside the network topological features. Incorporation of additional data like protein spatial locality into the PIN may furnish a more effective method and generate more reliable results. We establish our model’s performance on both blind and independent datasets. We also perform a detailed comparison of our model’s performance with the other existing methods and we observe that PINDeL outperforms the other methods. In addition, we apply the model to a set of experimentally reviewed proteins which is not used for training or testing and is not a part of the blind dataset. We then validate the PINDeL identified disease-proteins by functional annotation including Gene Ontology, disease, and KEGG pathway enrichment analysis. Our analysis on the host-pathogen PPIs informs that significant number of the PINDeL predicted proteins are found to be truly linked with diseases. Being an automatic classifier, PINDeL is useful for large-scale application in identifying the proteins associated with various genetic diseases in human. For public use, we make the executable file, and the detailed information available through GitHub repository at https://github.com/pralay-mitra/PINDeL, and at https://github.com/pralay-mitra/PINDeL/blob/main/Supplementary.xlsx, respectively.

## 2 Materials and Methods

### 2.1 Data set preparation

#### 2.1.1 Human PPI data

Following the methodology as described by Das et. al (Das et al., 2019), we construct Protein Locality Graph (PLG) of human consisting of 41550 proteins (represented as PLG nodes) and 8943744 PPIs (denoted as PLG edges) utilizing experimental knowledge from several online PPI resources including DIP (Salwinski et al., 2004), MINT (Licata et al., 2012), IntAct (Orchard et al., 2014), BioGRID (Oughtred et al., 2019), and pathway information from KEGG (Kanehisa et al., 2016) (Supplementary Section S1). We enrich the PLG with the maximum amount of known biological data for human by integrating information from Human Protein Reference Database (HPRD) and Online Predicted Human Interaction Database (OPHID) as mentioned below.

The HPRD (release 9, release time: April 13, 2010, access time: March 18, 2020) consists of human PPI data (81694 PPI among 12527 proteins) which is extracted from the literature by the expert biologists after thoroughly analyzing and interpreting the published data (Keshava Prasad et al., 2009). After extracting the interaction information from HPRD, we transform the PPI data into a graph by assigning edge weights denoting spatial locality scores to the interactions following the methodology proposed by Das et. al (Das et al., 2019).

The OPHID is a collection of human PPI data from several sources (Brown and Jurisica, 2005). OPHID includes literature-curated interactions from the Biomolecular Interaction Network Database (BIND) (Bader et al., 2003), HPRD (Keshava Prasad et al., 2009), and MINT (Licata et al., 2012). Interactions identified by high-throughput yeast two-hybrid mapping approach (Rual et al., 2005; Stelzl et al., 2005) are also inserted into the OPHID. Again, interolog-based interactions are predicted from *Saccharomyces cerevisiae, Caenorhabditis elegans, Drosophila melanogaster*, and *Mus musculus*. These are the potential interactions of human, which are predicted from the interactome data of the aforementioned model organisms given evolutionary conservation of the two known partners. We consider the human interactome from OPHID (Release time: March 3, 2020, Access time: March 18, 2020) that consists of 469515 PPIs and 18265 proteins. Next, we transform the OPHID PPI data into a weighted graph by assigning edge weights denoting spatial locality scores to the interactions following the methodology as proposed by Das et. al (Das et al., 2019).

Now, we incorporate human interaction information as extracted from the HPRD and OPHID into the PLG of human by taking union on the nodes and edges of PLG and the graphs constructed from HPRD and OPHID. Multiple edges between a pair of nodes will be replaced by a single edge with the corresponding weight value being set as a maximum of the variable edge weights carried by the multiple interactions. Total number of PPIs and unique proteins in the final PLG are 9128128 and 43919, respectively. The human Protein Locality Graph (PLG) includes the experimentally verified physical binary Protein-Protein Interactions (PPIs) from MINT, IntAct, DIP, and HPRD. Genetic interactions from BioGRID and the KEGG database are mapped to their corresponding PPIs and inserted into the PLG. The PLG also includes computationally predicted PPIs from the OPHID and the KEGG database. Hence, the human PPI data considered by our model include physical binary PPIs as well as computationally predicted PPIs. The topological features are mostly dependent on the connectivity of the network and majority of the features become inconvenient if the input graph is disconnected. Hence, we extract the largest connected component from the PLG of human and utilize it for the identification of the disease-associated proteins. Majority of the small sub-networks contain only a few nodes and edges, whereas, the largest connected network component (main component) of the PLG of human that is termed as the human PLG (*HPLG*) consists of 9128005 interactions and 43751 proteins.

#### 2.1.2 Disease-protein data

We collect disease-associated human genes from the DisGeNET database which integrates information of human Gene-Disease Associations (GDAs) and Variant-Disease Associations (VDAs) from several repositories (Piñero et al., 2016). The public repositories considered by DisGeNET include BeFree data (Bravo et al., 2015), Literature Human Gene Derived Network (LHGDN) (Bundschus et al., 2008), Genetic Association Database (GAD) (Becker et al., 2004), Mouse Genome Database (MGD) (Eppig et al., 2015), Rat Genome Database (RGD) (Shimoyama et al., 2015), Orphanet (Rath et al., 2012), ClinVar (Landrum et al., 2016), UniProtKB (Consortium, 2019b), Comparative Taxicogenomics Database (CTD) (Davis et al., 2015), and GWAS Catalog (Welter et al., 2014). DisGeNET is a comprehensive collection of expert-curated and text-mining derived disease-associated genes from the above mentioned public repositories and literature.

We download the curated gene-disease associations from the DisGeNET database (Release time: May 4, 2020, Access time: July 7, 2020) and extracted all the disease-associated genes. Total number of genes, diseases, and GDAs in DisGeNET are 9703, 11181, and 84038, respectively (Piñero et al., 2016). There exist 137822 literature evidences for these gene-disease associations listed in the database. Next, we utilize the mapping table provided by DisGeNET to map the gene identifiers to their corresponding UniProtKB identifiers. The UniProtKB ids of 542 genes are missing from the mapping table. As a result, we obtain 9161 human disease-proteins from the DisGeNET database. Among these 9161 proteins, 8887 proteins are present in the HPLG, and thus will be used for further analysis.

#### 2.1.3 Building training samples

The list of known disease-proteins, obtained from the DisGeNET database and mapped to the principal connected component of the human PLG, is called the ‘disease-protein set’ or the positive training dataset which consists of 8887 known disease-associated proteins. Assembling a list of genes or proteins that are known not to be involved in any hereditary disease is difficult or even impossible currently (Xu and Li, 2006). The genes present in the human genome can be classified into three categories - disease, non-disease, and essential genes. Genes associated with at least one disease are called disease-genes, whereas, rest are known as non-disease-genes. Alternatively, essential genes have features that differ significantly from both disease as well as non-disease-genes. Management of the essential genes as a unique group is beneficial for an effective comparison between the disease and non-disease genes (Tu et al., 2006). Also, proteins coded by the essential genes may not be responsible for any disease in human (Yu et al., 2020).

Since there exist no well-defined human essential genes, an approximated list of essential genes have been compiled by Tu et. al (Tu et al., 2006) and are also termed as the Ubiquitously Expressed Human Genes (UEHGs). A number of 1789 UEHGs have been reported and each UEHG is represented by its corresponding NCBI-UniGene identifier. We use the Database for Annotation, Visualization and Integrated Discovery (DAVID v6.8) for converting the UniGene ids of the UEHGs into their corresponding UniProtKB identifiers (Dennis et al., 2003). We get 2571 proteins after mapping the UniGene ids of the UEHGs to their corresponding UniProtKB ids by using the DAVID tool. Since 807 out of 2571 proteins are already being marked as disease-associated proteins by DisGeNET, so, we consider the rest i.e., 1764, as the essential protein group for this study. We can even call this group as the Ubiquitously Expressed Human Proteins (UEHPs). From the protein population present in the principal connected component of the human PLG and which are not listed in the DisGeNET, we exclude the UEHPs and then we call the remaining proteins as the ‘control-protein set’. Thus, the size of the ‘control-protein set’ is 33100 (43751 − (8887 + 1764)). From the ‘control-protein set’, we randomly select proteins with a size equal to that of the disease-proteins as the negative training dataset. Our final training data consisting of 8887 positive training samples and 8887 negative training samples is fed into the deep learning model.

### 2.2 Model construction

#### 2.2.1 Topological feature selection

We calculate 17 topological features for each of the *HPLG* proteins to train the classifier in distinguishing between a negative control protein node and a disease-related protein node. The features include Average Shortest Path Length (*ASPL*), Betweenness Centrality (*BC*), Weighted Betweenness Centrality (*WBC*), Closeness Centrality (*CC*), Clustering Coefficient (*CL*), Weighted Clustering Coefficient (*WCL*), Node Degree (*ND*), Weighted Node Degree (*WND*), Eccentricity (*E*), Average Neighbor Degree (*AND*), Weighted Average Neighbor Degree (*WAND*), Degree Centrality (*DC*), Information Centrality (*IC*), Weighted Information Centrality (*WIC*), Load Centrality (*LC*), Weighted Load Centrality (*WLC*), and Harmonic Centrality (*HC*). Details on the selected topological features are mentioned in the Supplementary Section S2. Features like *ASPL, BC, CC, CL, ND*, and *E*, have been previously used for the identification of the Alzheimer Disease-associated genes (Jamal et al., 2016) and other infectious disease-associated host genes (Barman et al., 2019).

We compute the Pearson Correlation Coefficients (PCCs) among the feature pairs to check the level of correlation among the features. Supplementary Figure S1 shows the heatmap representing the PCC values among the feature pairs. The pairwise correlation coefficients shown in the heatmap clearly indicate a significant level of correlation among a few sets of features. Identification and elimination of the features acting as proxies for another feature, will facilitate us in producing a robust and accurate model. Therefore, in order to avoid multicollinearity, we need to exclude the highly correlated variables from subsequent analysis. For this purpose, we initially standardize the features using StandardScaler of the Scikit-learn module (Pedregosa et al., 2011), which transforms the feature data such that the resulting distribution bears a mean value of 0 and standard deviation of 1. Finally, we perform Principal Component Analysis (PCA) on the standardized feature set to reduce the dimensionality of the original data set by generating a set of uncorrelated orthogonal Principal Components (PCs). We utilize these resulting PCs as the new features for our study and thereby, getting rid of the highly correlated features from further analysis.

#### 2.2.2 Classification

Identification of disease-associated human proteins is a binary classification problem where a particular protein is either associated with disease or not associated with any disease. We design a Graph Convolutional Network (GCN)-based (Kipf and Welling, 2016)) deep learning framework viz., *P*rotein *I*nteraction *N*etwork-based *De*ep *L*earning (PINDeL) for the identification of disease-associated human proteins. Figure 1 details the flow of our proposed PINDeL.

**Figure 1:**
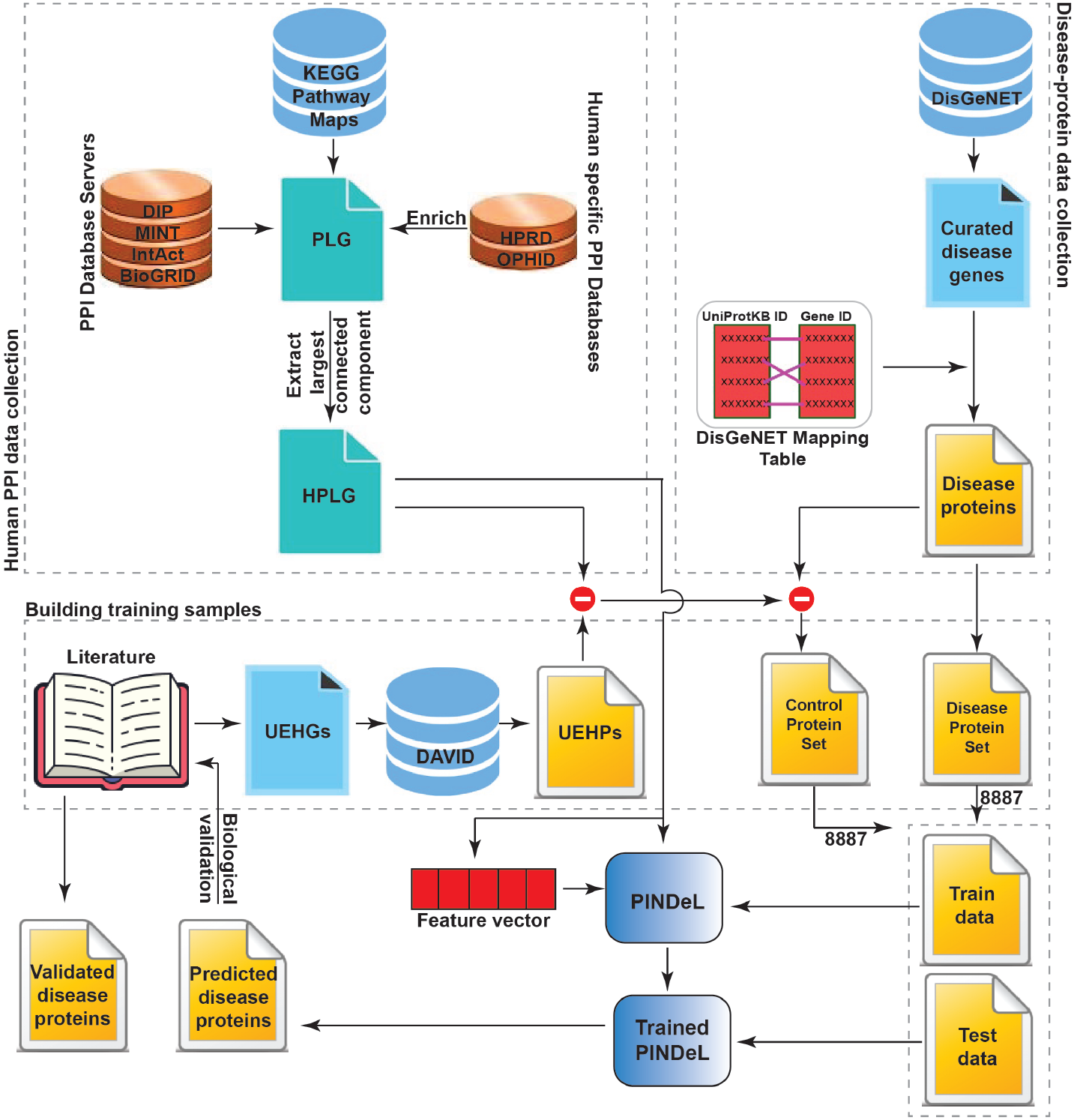
Flow diagram of the PINDeL describing the entire proposed methodology including collection of human PPI data, disease-protein data and generation of the HPLG and training samples, respectively, followed by the model training and testing procedures.

The GCN scales linearly in the number of graph edges and it can learn the hidden layer representations which encode both the local graph structure as well as the features of the nodes. GCNs are mainly designed for classifying nodes in a graph when labels are available for a small subset of nodes. The primary component of our designed model, PINDeL, is the GCN which can learn the embeddings from the node-specific information, nodes’ neighborhood, and the input network topology. Given a graph, *G*(*V, E*) with *N* nodes and *F* ^0^ input features for each node, a GCN takes an adjacency matrix *A* of *G* which is an *N* × *N* matrix representation of the graph structure, and a feature matrix *X* which is an *N* × *F* ^0^ matrix representation.

The propagation rule employed by GCN is

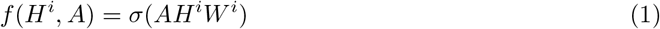

where *W* ^*i*^ is the weight matrix for layer *i* and *σ* is a non-linear activation function.

A hidden layer in GCN can be represented as *H*^*i*^ = *f* (*H*^*i*−1^, *A*), where *H*^0^ = *X* and *f* is the propagation rule as mentioned above. Each layer *H*^*i*^ corresponds to an *N* × *F* ^*i*^ feature matrix where each row is a feature representation of a node. At each layer, these features are aggregated to form the next layer’s features using the propagation rule *f* . Thus, at each consecutive layer, the features become more abstract. Mainly, GCNs vary according to different propagation rule representations (Kipf and Welling, 2016). Despite its simplicity, GCN is quite powerful.

But, there are two major limitations of the GCNs. Multiplication with *A* in Equation 1 means that, for every node, all the feature vectors of all the neighboring nodes are summed up, except the node itself. This is the first major limitation of GCN. Now, if there are self-loops in the input graph, then a particular node is also summed up along with the feature vectors of its neighboring nodes. But, when the input network does not possess self-loops, we need to add the identity matrix to *A* and then proceed with Equation 1, thereby fixing the first limitation (Kipf and Welling, 2016). The second major limitation is that, *A* is typically not normalized and therefore the multiplication with *A* (Equation 1) will completely change the scale of the feature vectors. To get rid of this problem, we need to normalize *A* such that all rows sum to one, i.e. *D*^−1^*A*, where *D* is the diagonal node degree matrix. Multiplying with *D*^−1^*A* corresponds to taking the average of neighboring node features. Hence, we need to perform a symmetric normalization, i.e. 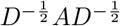, for not staying limited to taking the average of only the neighboring node features (Kipf and Welling, 2016). Combining the solutions of the two major limitations of the GCNs, we get the modified propagation rule which is

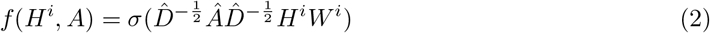

with *Â* = *A* + *I*, where *I* is the identity matrix and 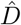 is the diagonal node degree matrix of *Â*.

Our model, PINDeL, is composed of two graph convolutional layers, each followed by a nonlinear Rectified Linear Unit (ReLU) activation function and a Dropout operation (Figure 2). Transformation of the summed weighted input from the node into the activation of the node is carried out by the activation function in a neural network. The ReLU activation function is a piecewise linear function that will output the input directly if it is positive, otherwise, it will output zero. Dropout operation refers to ignoring both hidden and visible neurons in a neural network and is mainly applied to prevent overfitting. After the second graph convolutional layer, two fully-connected layers are applied. Again, to overcome overfitting, a dropout operation is employed after the first fully-connected layer. The classification output layer has two classes whose activation is regulated by a Softmax function. The input values, which can be positive, negative, zero, or greater than one, are mapped into a set of values in between 0 and 1 by the Softmax function in order to interpret the values as probabilities. PINDeL utilizes the cross-entropy loss function also known as the logistic loss or the log loss. It measures the performance of a classification model whose output is a probability value lying between 0 and 1. Cross-entropy loss increases as the predicted probability diverges from the actual label. For training the neural network, we use the Adam (Adaptive Moment Estimation) optimizer for its speed and rapid convergence, although it is computationally expensive.

**Figure 2:**
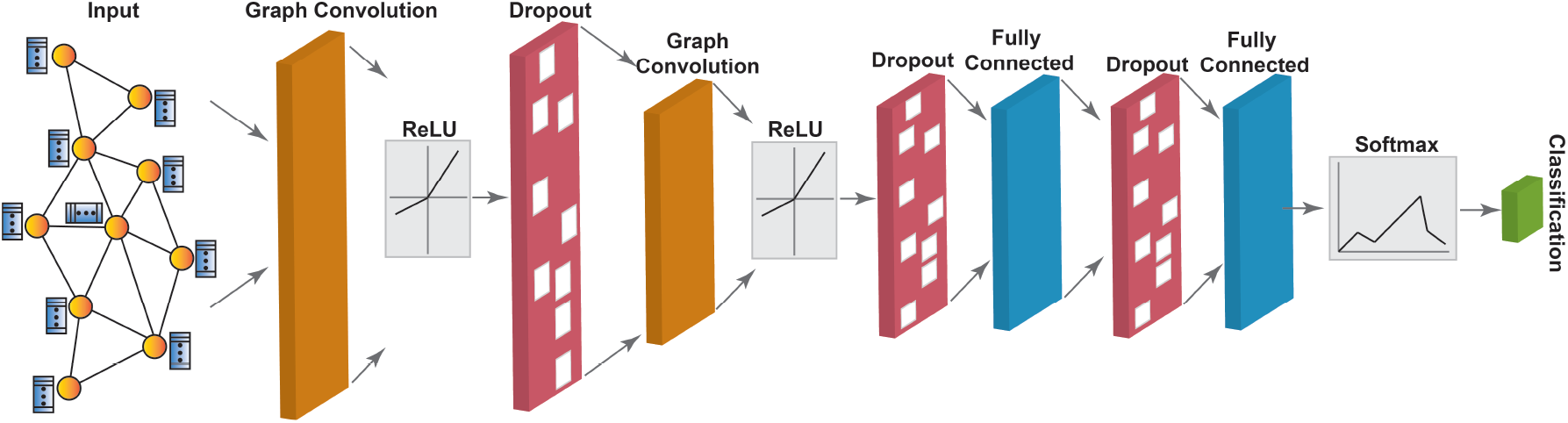
Structure of PINDeL model. The input of our model contains two components, HPLG and additional information for the nodes. The gradient colored input nodes represent proteins and the connections between the nodes represent protein-protein interactions. The vectors shown alongside the nodes represent the node feature vectors. PINDeL, is composed of two graph convolutional layers, each followed by a nonlinear ReLU activation function and a Dropout operation. The graph convolutional layers are used for learning the node embeddings. For each node, the GCNs aggregate information from the previous layer embeddings of its neighboring nodes. Two fully connected layers follow the GCNs. Dropout is applied to overcome overfitting. The classification output layer is regulated by a binary Softmax function.

#### 2.2.3 Model hyperparameters

We optimize the performance measures of the prediction model by performing grid search-based hyperparameter tuning. We also apply grid search-based technique for optimizing the architecture of the model. Supplementary Table S1 illustrates briefly the selection of the best hyperparameters. In terms of implementation, we incorporate two GCN layers in our model, with the dimension of the hidden representation as 512 and the final embedding dimension as 256. Following the GCN layers, we set the number of fully connected dense layers as two, with the dimension of the hidden representation as 256 and the final classification dimension as 2. For training the model, we use Adam optimizer with the learning rate as 0.001. To reduce overfitting, we apply a combination of dropout layers to the hidden layers with the dropout rate as 0.5 along with the weight decay method.

## 3 Result and Discussions

### 3.1 Performance of PINDeL

Firstly, we assess the goodness of our model in detecting disease-associated proteins from the DisGeNET database using the 10-fold cross-validation strategy when PLG of human and 17 node-based topological features are supplied as inputs. We distribute the entire dataset into 10 segments or folds of equal (or nearly equal) sizes. We repeat training and testing 100 times, where in each time we randomly prepare 9 sets or folds being used for training, whereas, the remaining one set or fold being utilized for testing. The average performance of the classifier over 10 folds provides a measure of the overall performance of the model. Measures of the evaluation metrics, as discussed in Supplementary Section S3, obtained during 10-fold cross validation, are summarized in Figure 3.

**Figure 3:**
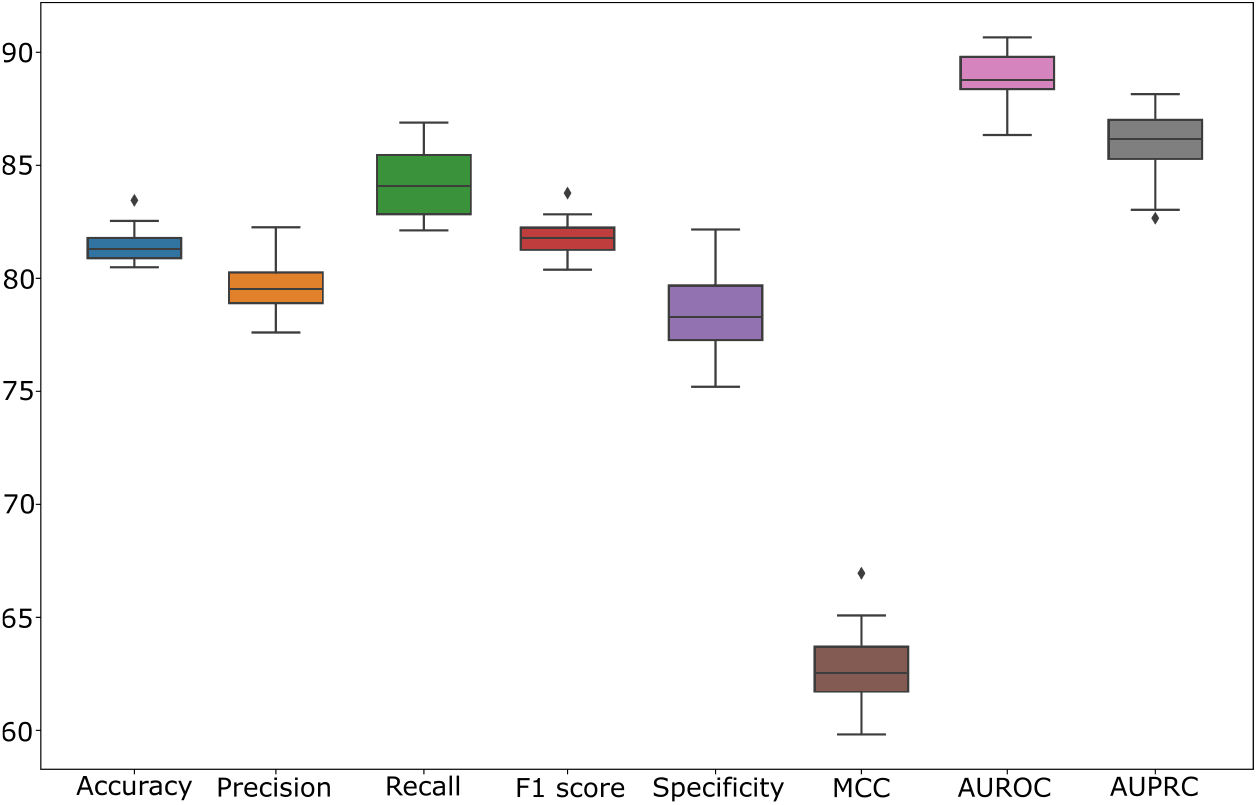
Statistical results of 10-fold cross validation.

The average values (standard deviation) of accuracy, precision, recall, F1-score, specificity, MCC, AUROC, and AUPRC are 81.42% (±0.532), 79.67% (±0.81), 84.13% (±1.55), 81.82% (±0.59), 78.54% (±0.32), 62.79% (±0.52), 88.90% (±0.92), and 86.05% (±1.24), respectively. The accuracy value indicates that the classifier correctly predict 81.42% of the total predictions. Here, we select the negative training samples from the control-protein set that may include disease-associated proteins, but have not been experimentally verified yet. Therefore, PINDeL may classify some negatively labeled proteins as disease-associated proteins that leads towards an increase in the number of the false positives and a low precision score. Hence, for this disease-protein classification problem, recall measure will be a better metric than precision. A good predictive performance of the model will be reflected by less number of false negatives and thereby a higher recall score, which in our case is 84.13%. High F1-score (81.82%), AUROC (88.90%), and AUPRC (86.05%) indicates commendable classification performance. The highest training, validation, and testing accuracy achieved during 10-fold cross-validation is 83.18%, 83.50%, and 83.45%, respectively. The AUROC value of 0.90 (Figure S2(a)) indicates a significantly good predictive capability of the model in discriminating between the positive and the negative results. The AUPRC value of 0.88 (Figure S2(b)) indicates high precision relating to a low false positive rate and high recall relating to a low false negative rate. We choose the model which shows the highest AUROC value as the final prediction model. To reduce the risk of degree data bias, we split the proteins into hub proteins and non-hub proteins according to their degrees and feed them to PINDeL. We noted that, for the non-hub (hub) proteins, the accuracy, precision, recall, F1-score, specificity, MCC, AUROC, and AUPRC are 80.65%(80.80%), 75.80%(76.03%), 88.21%(88.24%), 81.54%(81.68%), 73.55%(73.79%), 62.23%(62.50%), 89.12%(89.28%), and 86.48%(86.73%), respectively.

The applicability and the robustness of our method is also tested for the real-world situations, where the size of the negative dataset is often much larger than that of the positive dataset. On such an imbalanced dataset where the size of the negative samples is twice that of the positive dataset, the accuracy, precision, recall, F1 score, specificity, MCC, AUROC, and AUPRC of PINDel is 76.96%, 89.22%, 61.28%, 72.65%, 92.61%, 56.75%, 90.75%, and 88.70%, respectively. We can observe that, with an increase in the size of the negative dataset, the specificity increases. However, with an increase in the size of the negative dataset, recall decreases. Undoubtedly, most of the learning models including our model performs better with balanced dataset compared to the imbalanced dataset.

### 3.2 Comparison with the existing works

Several existing works have considered the human PIN as a primary input for determining the disease-associated genes or proteins. Also, a sizeable amount of data like protein sequence features, PIN features, disease symptoms, shortest paths, is needed by the existing research studies to train their corresponding models. Our method utilizes a PIN-type graph called the Protein Locality Graph as primary input that includes the protein spatial locality as an additional data in the form of interaction/edge weights denoting the locality scores. Higher the weight of the edge/interaction connecting two proteins, lesser is the distance between them. Utilizing only the PLG and the 17 topological features for training, our framework performs significantly well.

Again, machine learning-based techniques including SVM, NB, RF, and deep learning-based techniques including CNNs and image-based CNNs have been utilized for this specific research problem. Although, the GCNs operate similarly like the CNNs, but there exists a major difference between them. CNNs are designed specially to operate on Euclidean or regular structured data, whereas the GCNs, being a generalized version of the CNNs, operate on irregular or non-Euclidean structured data. Thus, GCNs can be especially applied to model real-world entities such as graphs. Therefore, for the biological networks such as PINs, GCN-based models will be significantly effective as that compared to traditional CNNs or image-based representations of PIN and disease data.

We perform a thorough comparison between our GCN-based PINDeL and the other existing machine-learning and deep learning-based techniques of disease gene/protein identification in human. It should be noted that there exist significant differences between the datasets and the features utilized for training the corresponding models of all the methods. Xu et al. (Xu and Li, 2006) trained their KNN classifier on the human PPI dataset derived from the Online Predicted Human Interaction Database (OPID). This method utilizes five topological features including degree, 1N index, 2N index, average distance to disease genes, and positive topology coefficient. The DNN model proposed in (Barman et al., 2019) exploits the expert-curated human PPIs derived from the Human Protein Reference Database (HPRD). This study uses nine topological features and protein sequence features including Amino Acid Composition (AAC), Dipeptide Composition (DC), Pseudo-Amino Acid Composition (PAAC), and Conjoint Triad Descriptors (CTD). Since the evaluation metric measures have been reported in (Barman et al., 2019) for different feature combinations, we compute the average of all those values and report it in Table 1. The RF classifier (Yu et al., 2020) has been trained on the human interactome datasets derived from I2D database based on its corresponding 14 topological features. The input dataset in our case is the HPLG, a graph we construct by integrating both PPI and protein locality information. The other methods (Xu and Li, 2006; Barman et al., 2019; Yu et al., 2020) consider only the protein interaction data as the input dataset. Whereas, HPLG is more advanced and contains more information because apart from including the PPI data like others, it also incorporates the protein spatial locality information in the form of edge weights or confidence scores. We train our GCN-based model, PINDeL, utilizing 17 topological node-based features of the HPLG. Table 1 indicates that not all the existing methods did a systematic analysis on all the evaluation metric (not available data is marked as NA). Nevertheless, Table 1 demonstrates that PINDeL outperforms the other existing methods in accuracy, recall, F1-score, specificity, AUROC, and AUPRC. Only RF classifier (Yu et al., 2020) shows better precision compared to PINDeL, but its recall measure is quite low. For disease gene/protein identification problem, recall is a more suitable and better evaluation metric than precision.

**Table 1:** The overall performance of PINDeL compared to existing methods. Not available information is marked as NA.

| Method | Accuracy (%) | Precision (%) | Recall (%) | F1-score (%) | Specificity (%) | AUROC (%) | AUPRC (%) |
| --- | --- | --- | --- | --- | --- | --- | --- |
| **GCN-based PINDeL** | **83.45** | **80.50** | **86.89** | **82.83** | **79.89** | **90.66** | **88.15** |
| RF (Yu et al., 2020) | 77 | 88 | 77 | NA | NA | 88 | NA |
| DNN (Barman et al., 2019) | 78.62 | NA | 83.89 | 80.16 | 73.03 | 83.31 | NA |
| KNN (Xu and Li, 2006) | 76 | 76 | 75 | NA | NA | NA | NA |

For a detailed extensive comparison, we downloaded the data of (Yu et al., 2020; Xu and Li, 2006) from the I2D database (Interologous Interaction Database) (Brown and Jurisica, 2007) also known as the OPHID database (Brown and Jurisica, 2005), which consists of 18, 265 proteins and 458, 160 PPIs. Now, the inputs required by our model include a weighted graph called the Protein Locality Graph (PLG) and its corresponding node-based 17 topological features. The PLG is a Protein Interaction Network (PIN) whose edge weights denote a spatial locality score computed using the network-based zoning approach (Das et al., 2019). We add unit edge weights to the PIN constructed from I2D, so that it will be compatible with the GCN-trained model, PINDeL. Further, we extract the largest connected component (18, 185 proteins and 458, 136 PPIs) from this I2D-based PIN, which is an utmost necessity for a valid computation of the 17 topological features. Then, we feed this data to our trained model. For this I2D data utilized by Yu et. al. (Yu et al., 2020) and Xu et. al. (Xu and Li, 2006), the values of accuracy, precision, recall, F1-score, specificity, MCC, AUROC, and AUPRC are 55.85%, 51.84%, 86.25%, 64.76%, 28.83%, 18.25%, 64.22%, 59.65%, respectively. For the data utilized by Barman et. al. (Barman et al., 2019), the average values of accuracy, precision, recall, F1-score, specificity, MCC, AUROC, and AUPRC are 55.37%, 71.41%, 60.43%, 65.46%, 43.54%, 37.12%, 53.97%, 73.24%, respectively. We also apply the existing traditional machine learning approaches on our data including logistic regression, random forest, SVM, Naive-Bayes, decision tree, KNN, and AdaBoosting. The 10-fold cross validation performance metrics of these approaches as compared to our PINDeL model are clearly detailed in Table 2. From Table 2, we can see that although the accuracy of the model trained by AdaBoosting is the highest among all the methods, its recall is quite low which must be significantly high for a good predictive model performance. Similarly, although the AUROC measure of RF is quite high, but its other performance metrics are less than PINDeL. Therefore, based on the statistics shown in Table 2, we can safely conclude that PINDeL provides a significantly good predictive performance as that compared to the other existing traditional machine-learning approaches.

**Table 2:**
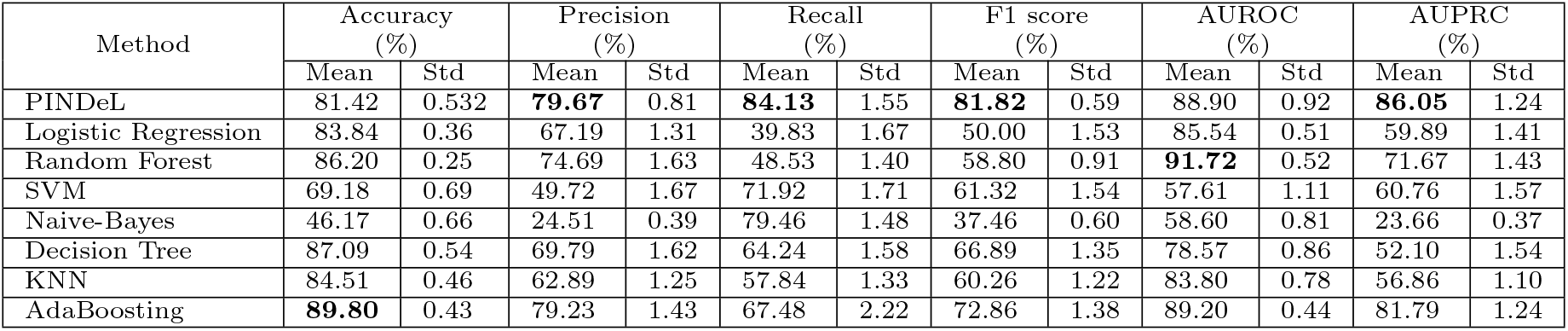
The overall performance of PINDeL compared to the existing methods.

### 3.3 Application on novel disease-protein

#### 3.3.1 Predicting novel disease-proteins

We predict infectious diseases-associated proteins using PINDeL from an independent dataset consisting of the reviewed human proteins which are not used for training or testing purposes. Firstly, we include into the independent dataset, 100 proteins which are not used for training or testing purposes but are associated with at least one disease as per the DisGeNET database. By definition, our control proteins are not associated with any disease as per DisGeNET. Excluding all the control proteins that we utilize for model learning, we feed the remaining 24220 control proteins into our independent dataset. Thus, our independent dataset consists of a total number of 24320 proteins. We apply PINDeL to this dataset for identifying the disease-associated proteins from it. We find that PINDeL recognizes 94 proteins out of the 100 proteins belonging to the curated disease-protein set of the DisGeNET database. Among the 24220 control proteins, PINDeL marks 6354 proteins as disease-proteins. Altogether, PINDeL identifies 6448 disease-proteins among 24320 proteins present in the independent dataset. Although PINDeL classifies 94 proteins are already a part of the DisGeNET database, but the 6354 proteins are not part of DisGeNET, and accordingly, we initially marked them as control set. But PINDeL identifies these 6354 proteins as disease proteins. Our detailed analysis indicates that 3196 out of 6448 disease-proteins are validated either through PHISTO (Pathogen-Host Interaction Search Tool) or using enrichment analysis. Interestingly, 573 diseases-proteins are validated by PHISTO and by enrichment analysis both (Figure 4). Now, we proceed with a thorough biological validation of these 3196 putative PINDeL predicted disease-proteins.

**Figure 4:**
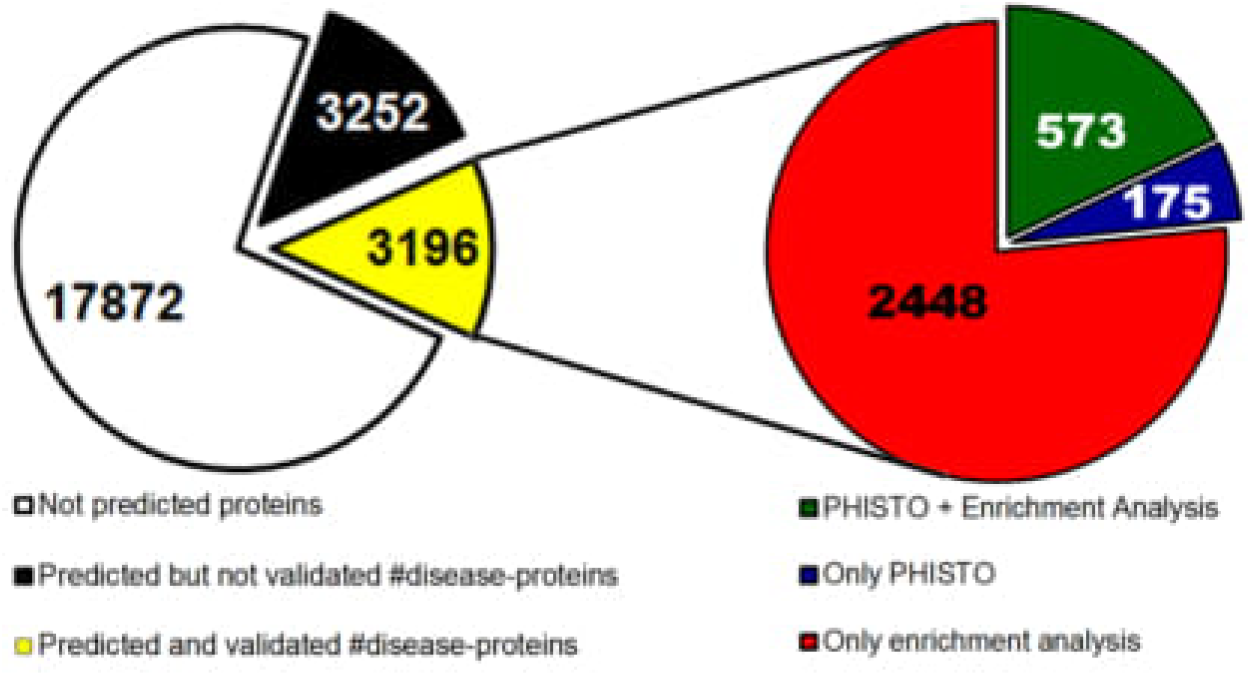
Summary of validation of the novel disease-associated proteins identified by PINDeL.

#### 3.3.2 PHISTO-based validation

PHISTO is the most comprehensive experimentally-verified pathogen-human protein-protein interaction database for studying infection mechanisms through Pathogen-Host Interactions (PHIs) (DurmuÇ Tekir et al., 2013). PHISTO provides us with access to the most up-to-date PHI data for all pathogen types which are experimentally verified protein interactions with hosts (DurmuÇ Tekir et al., 2013). For validation and functional annotation of the PINDeL identified disease-proteins, we use PHISTO (access on: August 09, 2020) that includes data for 588 pathogen strains, 48615 PHIs, 4455 targeting pathogen proteins, and 8373 targeted human proteins. Among the 3196 putative disease-proteins identified by PINDeL, 748 proteins belong to PHISTO (Figure 4) and hence, are valid targeted human proteins associated with one or multiple infectious diseases. Among the 100 DisGeNET disease-proteins, PINDeL identifies 94 as disease-associated of which only 46 are present in PHISTO. Hence, it informs that different existing disease databases are not in synchronization with each other with respect to disease-protein related information for human. However, among the 24220 control proteins, which are claimed not to be associated with any disease by DisGeNET, PINDeL identifies and PHISTO validates 702 proteins that belongs to disease-associated proteins in human (Figure 4).

#### 3.3.3 Enrichment analysis

Gene Ontology (GO) hierarchically classifies the functions of the genes into terms organized as a graphical structure or as an ontology. Such terms are grouped into three categories - Biological Process (BP), Molecular Function (MF), and Cellular Component (CC) Consortium (2019a). Biological Process describes the larger cellular or physiological roles executed by the genes. Molecular Function denotes the molecular activity of a gene. Cellular Component denotes the cellular location where the gene executes its function. Interpreting sets of genes by utilizing the Gene Ontology classification system is known as GO term enrichment. For a set of genes that are up-regulated under certain conditions, a detailed GO enrichment analysis determines which GO terms are over-represented or under-represented by exploiting the annotations for that gene set. Gene Ontology enrichment analysis shows that the genes corresponding to 3305 (2142 unique) putative disease-associated proteins are enriched in biological processes like viral process (GO:0016032), regulation of tumor necrosis factor-mediated signaling pathway (GO:0010803), positive regulation of phosphorylation (GO:0042327), protein autophosphorylation (GO:0046777), intracellular transport of virus (GO:0075733), etc. Figure 5(A) shows the significantly enriched GO biological process terms for the PINDeL identified disease-associated proteins and the relationship between the enriched biological processes. Two biological processes (nodes) are connected if they share 20% (default) or more genes. Darker nodes represent more significantly enriched gene sets. Bigger nodes represent larger gene sets and thicker edges represent more overlapped genes (Ge et al., 2020). In Figure 5(A), we show the top 50 most significantly enriched GO terms generated by setting a threshold of 0.05 on the P-value. KEGG pathway enrichment analysis, performed using ShinyGO (Ge et al., 2020), shows that 2894 putative disease-associated proteins are enriched in KEGG pathways like necroptosis, metabolic pathways, olfactory transduction, etc as shown in Figure 5(B). It is noteworthy that 489 predicted disease-proteins participate in Herpes simplex virus 1 infection pathway.

**Figure 5:**
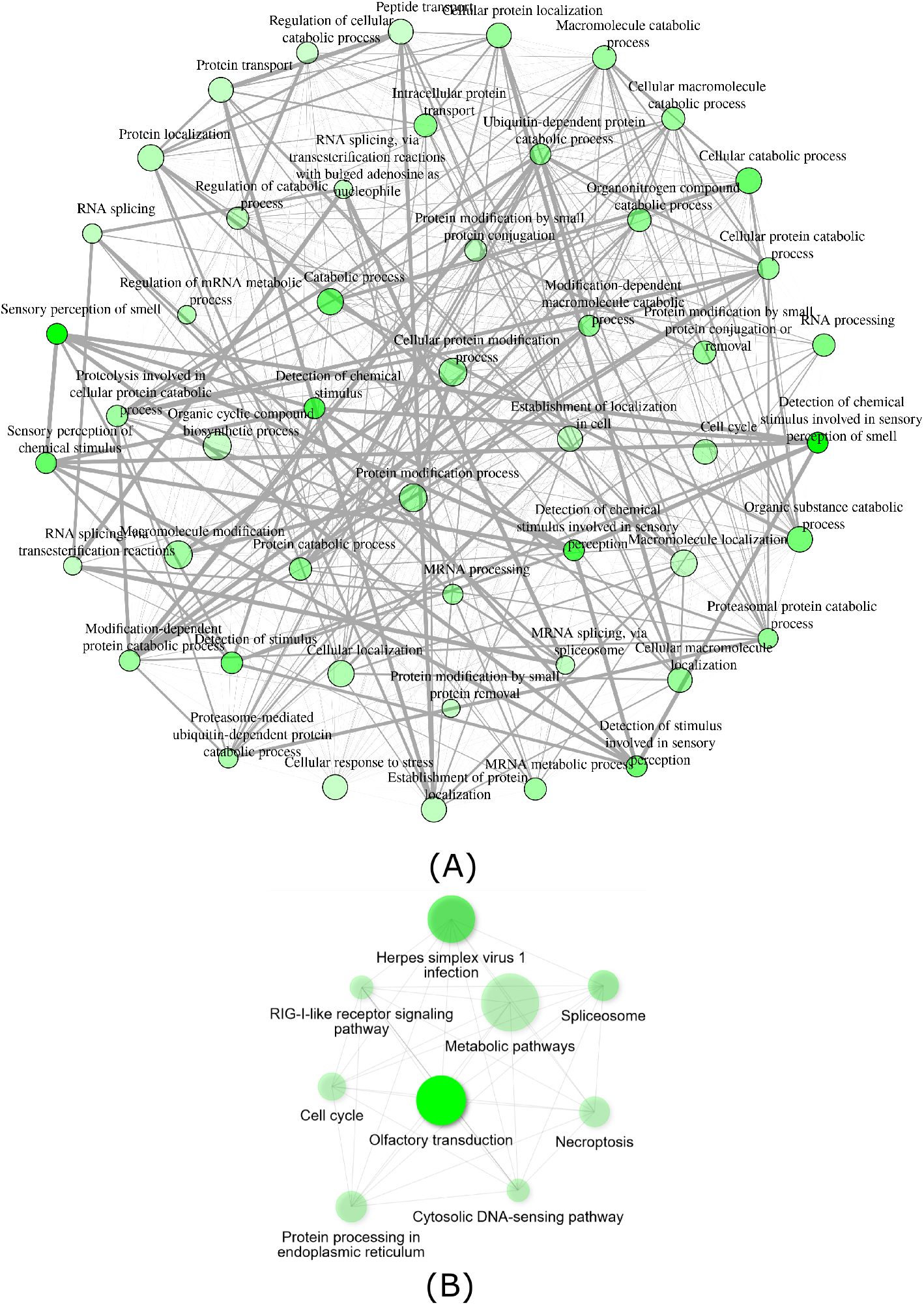
(A) Significantly enriched Gene Ontology Biological Process terms corresponding to the putative disease-associated proteins identified by PINDeL. For the visualization, we use graphical gene-set enrichment tool ShinyGO (Ge et al., 2020). (B) Significantly enriched KEGG Pathway terms corresponding to the putative disease-associated proteins identified by PINDeL.

#### 3.3.4 Annotation of the predictions

We functionally annotate PINDeL predicted 6448 putative disease-proteins using the Database for Annotation, Visualization and Integrated Discovery (DAVID) (Sherman et al., 2009; Huang et al., 2009) - a comprehensive set of functional annotation tools. The Genetic Association Database (GAD) is a database of genetic association data from complex diseases and disorders (Becker et al., 2004). GAD consists of a collection of disease classes along with the information about the genes belonging to one or multiple disease classes. Firstly, we utilize DAVID to determine the list of proteins from the putative predicted set, which belong to one or multiple GAD disease classes. Disease ontology enrichment analysis shows that 598, 713, 526, 419, 110, 602, and 188 proteins out of 3196 proteins are classified as disease terms, i.e., immune, chemdependency, pharmacogenomic, infection, normalvariation, cancer, and aging, respectively. In total, 1853 out of 3196 predicted proteins are found to be associated with one or multiple GAD disease classes as per DAVID. In addition, we observe that the highly predicted infectious disease-associated proteins are also found in cancer and immune disease terms. Similar to GAD, Online Mendelian Inheritance in Man (OMIM) database is a continuously updated catalog of human genes and genetic disorders and traits Amberger et al. (2015). Here, we also perform a disease ontology enrichment based on the OMIM database. It is noteworthy that 7, 6, 4, and 3 putative proteins are found to be associated with breast cancer, colorectal cancer, gastric cancer, and ovarian cancer, respectively. Detailed information is available at https://github.com/pralay-mitra/PINDeL/blob/main/Supplementary.xlsx.

## 4 Conclusion

Identification of disease-associated proteins is vital for exploring the underlying mechanisms and for combating infectious diseases. Although experimental techniques are the best candidates for addressing this problem, but computational methods are more effective in terms of cost and human resources. In addition, with an increasing availability of experimental data, the computational identification of disease-associated proteins becomes more accurate and easy.

Here, we present a novel Graph Convolutional Network-based deep learning framework called PINDeL which is capable of identifying the disease-associated proteins in human. We provide the Protein Locality Graph of human along with the node-based topological features of the graph as inputs to the model. The model achieves an AUROC value of 0.90 and an AUPRC value of 0.88, indicating the model’s excellent capability of identifying disease-associated proteins from the human PLG. We observe that balanced dataset perform better than the imbalanced datasets, which is an ideal case for any prediction model. From a detailed comparative analysis, we observe that PINDeL outperforms other existing disease-gene/protein identification models.

PINDeL is the first GCN-based deep learning framework for identifying disease-associated human proteins. The goodness of PINDeL is tested using 10-fold cross validation, whereas, the applicability and usefulness is verified on an independent dataset consisting of reviewed human proteins not used for training or testing purposes. Among 24320 proteins present in the independent dataset, PINDeL identifies 6448 putative disease-associated proteins. We find that 748 predicted disease-proteins are part of PHISTO, and hence are valid disease-associated proteins. Again, we perform a detailed disease ontology enrichment based on GAD where we find that 3067 (1853 unique) proteins are significantly enriched with disease terms like immune, infection, cancer, etc (Figure 6). Gene Ontology enrichment analysis shows that the genes corresponding to 3305 (2142 unique) putative disease-associated proteins are enriched in biological processes like viral process (GO:0016032), regulation of tumor necrosis factor-mediated signaling pathway (GO:0010803), positive regulation of phosphorylation (GO:0042327), protein autophosphorylation (GO:0046777), intracellular transport of virus (GO:0075733), etc (Figure 5(A)). KEGG pathway enrichment analysis shows that 2894 putative disease-associated proteins are enriched in pathways like necroptosis, metabolic pathways, olfactory transduction, Herpes simplex virus 1 infection pathway, etc (Figure 5(B)). These validation results suggest that the predictive power of the proposed model is quite good. Therefore, the remaining predicted protein set after validation still stays as highly probable disease-associated proteins which may be further presented for experimental validation.

**Figure 6:**
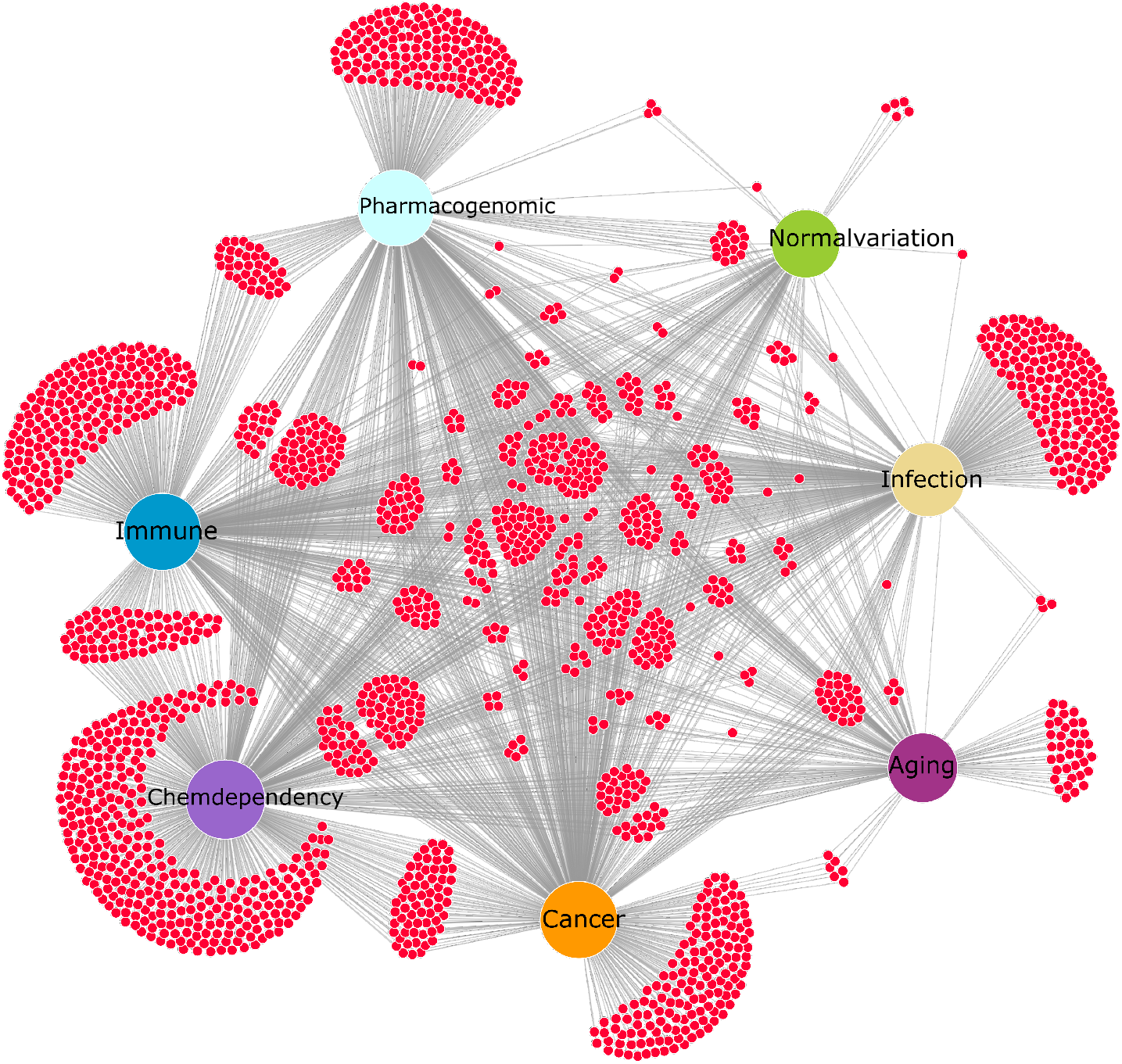
Overall representation of the functional annotation of the PINDeL identified putative proteins by DAVID based on GAD disease class by utilizing the DiVenn tool (Sun et al., 2019). Overlapping disease-proteins are shared by multiple GAD classes and there are 22 PINDeL identified novel disease-proteins which are associated with all the seven GAD classes including immune, infection, cancer, aging, chemdependency, normalvariation, and pharmacogenomic.

The 10-fold cross-validation performance metrics of PINDeL, as detailed in Result and Discussions section, reflect the good performance of the model. Computational identification of novel disease-associated proteins has become easier and more accurate with incorporating data heterogeneity in the training datasets, which is one of the limitations of our approach. Another limitation of traditional machine learning and deep learning approaches is the black box model concept, which limits human understanding of how various model parameters or variables are combined to generate the predictions. More detailed analysis and understanding is required to solve this black box concept. In future, it is also required to exploit GCN-based models utilizing homogeneous training datasets.

## Acknowledgement and Funding

This work was supported by the Open Competitive Grand Challenge Seed Grants (SGIGC) of Indian Institute of Technology Kharagpur (SRIC project code: WBC). B. D. is supported by an INSPIRE Fellowship (INSPIRE Code: IF150632) sponsored by the Department of Science and Technology, Government of India.

## Conflict of interest

There is no conflict of interest in the publication of this research study.

## Notes

### Competing Interest Statement

The authors have declared no competing interest.

